# Brain infiltrating T cells mediate microglial dysregulation and neuronal loss following SAH

**DOI:** 10.1101/2025.10.10.681332

**Authors:** F Moro, E Mazzone, R Pascente, M Bataclan, S Monticelli, F Pischiutta, MC Trolese, C Vasco, J Geginat, M Di Feliciantonio, F Ortolano, T Zoerle, ER Zanier

**Affiliations:** Laboratory of Traumatic Brain Injury and Neuroprotection, Department of Acute Brain and Cardiovascular Injury, Mario Negri Institute for Pharmacological Research IRCCS, Milan, Italy; Institute for Research in Biomedicine (IRB), Università della Svizzera italiana, Bellinzona, Switzerland; National Institute for Molecular Genetics “Romeo ed Enrica Invernizzi” INGM, Milan, Italy; Neuroscience Intensive Care Unit, Department of Anesthesia and Critical Care, Fondazione IRCCS Cà Granda Ospedale Maggiore Policlinico, Milan, Italy; Department of Pathophysiology and Transplantation, University of Milan, Milan, Italy

**Author notes:** Corresponding author contact information: Elisa R Zanier, Department of Acute Brain and Cardiovascular Injury, Istituto di Ricerche Farmacologiche Mario Negri IRCCS, Milan, Italy. Equal contribution first-author (F Moro, E Mazzone).

**Keywords:** subarachnoid hemorrhage, neuroinflammation, T cell, microglia

## Abstract

The contribution of T cells to neuroinflammation after aneurysmal subarachnoid hemorrhage (SAH) remains poorly understood. Using a murine pre-chiasmatic injection model of SAH we demonstrate that T cell infiltration into the brain modulates microglial activation and promotes neuronal death. Targeted transcriptomic profiling revealed a sustained neuroimmune response at 7 days post injury (dpi) characterized by a major involvement of T cells and microglia activation. Immunohistochemistry confirmed focal CD3+ T cell infiltration, predominantly CD4+, in the brain at the site of blood injection (BI), choroid plexus and meninges in SAH mice at 3- and 7-dpi. This temporal pattern was also observed in the CSF of a human SAH cohort. T cell presence spatially correlated with regions of microglial reactivity and neuronal loss. Notably, CD3-knockout mice exhibited reduced microglial activation and preserved neuronal viability. These findings identify T cells as key amplifiers of post-SAH neuroinflammation and neuronal damage. Targeting T cell–microglia crosstalk may represent a novel therapeutic avenue for SAH.

**Summary statement:** This study shows that brain T-cell infiltration after subarachnoid hemorrhage drives microglial activation and neuronal loss in mice, with similar patterns observed in patients. Data indicate T cells as key mediators of post-injury neuroinflammation with therapeutic implications.

## Introduction

Aneurysmal subarachnoid hemorrhage (SAH) is a rare but devastating form of hemorrhagic stroke affecting 10/100.000 individuals each year (Macdonald and Schweizer, 2017). SAH has a mortality rate of ∼50%, and survivors frequently experience persistent cognitive and functional impairments (Van Gijn, Kerr and Rinkel, 2007; Al-Khindi, Macdonald and Schweizer, 2010). Despite advancements in clinical management, SAH remains a disease with a significant morbidity, underscoring the urgent need to better understand its complex pathophysiology and identify therapeutic targets (De Oliveira Manoel *et al*., 2016; Thilak *et al*., 2024). After SAH, global cerebral ischemia initiates a cascade of pathophysiological processes collectively referred to as early brain injury (EBI) which occurs within the first 72 hours (Sehba *et al*., 2012; Chen *et al*., 2014). EBI is characterized by alteration of ionic homeostasis, microvascular dysfunction, blood brain barrier (BBB) breakdown, cerebral edema, neuroinflammation and oxidative stress, ultimately contributing to neuronal cell death (Lauzier *et al*., 2023). In the subacute phase patients are at risk of delayed cerebral ischemia (DCI) typically occurring between 3 and 14 days after the initial bleeding (Macdonald, 2014). While cerebral vasospasm (CVS) has been closely associated with DCI (Crowley *et al*., 2011), evidence suggests that additional mechanisms, such as micro thrombosis, cortical spreading depolarizations microcirculatory failure and neuroinflammation also play a critical role (Macdonald, 2014).

Secondary injury mechanisms, including the accumulation of blood breakdown products act as potent triggers for central nervous system (CNS) immune activation. This leads to the recruitment of both innate and adaptive immune cells into the brain amplifying local inflammation and potentially exacerbating tissue injury processes (Roa *et al*., 2020; Zhang *et al*., 2023). While the role of innate immune cells such as microglia and neutrophils has been explored, the involvement of adaptive immune responses, particularly T lymphocytes (T cells), remains poorly characterized in the context of SAH (Clatterbuck *et al*., 2003; Provencio *et al*., 2011; Hanafy, 2013; Provencio, 2013; Schallner *et al*., 2015; Schneider *et al*., 2015; De Oliveira Manoel and Macdonald, 2018). T cells are key orchestrator of adaptive immune response in the injured brain, capable of exhibiting toxic or regenerative properties depending on their phenotype, and the type and timing of injury (Kleinschnitz *et al*., 2010). In models of ischemic stroke and traumatic brain injury, CD4+ T cells have been shown to adopt proinflammatory phenotypes (Fee *et al*., 2003; Shi *et al*., 2023; Jiang *et al*., 2024), infiltrate brain parenchyma, and worsen tissue damage (Gate *et al*., 2021; Williams *et al*., 2021). Interestingly, emerging evidence suggest that activated T cells can modulate microglial behavior driving their transitions from homeostatic to disease associated phenotypes and amplifying the local inflammatory milieu (Benakis *et al*., 2022). However, a substantial knowledge gap persists in understanding the contribution of T cell response in injury evolution after SAH. While few observational studies have reported dynamic changes in CD4+ T cells subsets in peripheral blood and in cerebrospinal fluid (CSF) of SAH patients (Moraes *et al*., 2020; Chaudhry *et al*., 2021; Song *et al*., 2023), direct evidence of their infiltration into brain tissue and their impact on resident immune cells and neurons is lacking.

In this study, we aimed to elucidate the role of T cells in modulating neuroinflammation and neuronal injury following SAH. Using a murine pre-chiasmatic injection model, we investigated the spatial and temporal dynamics of T cell infiltration into the brain, their interaction with microglia, and their contribution to neuronal loss. In SAH patients, we analyzed T cell temporal changes in the blood and CSF, to assess whether centrally infiltrating T cells also represent a key pathological hallmark in human pathology. Our findings provide novel insights into T cell–mediated mechanisms of secondary brain injury after SAH and suggest that targeting T cell–microglia interactions may represent a promising therapeutic strategy.

## Results

### SAH induces vasospasm, cerebral edema and sensorimotor deficits

Induction of SAH led to an immediate and marked (>80% from baseline), drop in cerebral blood flow (CBF) as measured by laser speckle contrast imaging. This reduction persisted for 30 minutes post injury (p<0.001 at 0 min, 1 min, 5 min, p<0.01 at 30 min). Notably, this prolonged reduction in CBF was specifically attributable to subarachnoid blood, as CBF reduction in saline injected controls was significantly lower with faster recovery to baseline levels compared to SAH mice (p<0.001 1 min, 5 min, 30 min, SAH vs saline) (Fig. 1C). CBF drop in SAH mice was associated with acute neurological deficits as shown by increased righting reflex latency (p<0.001, compared to sham and saline controls) (Fig. 1D). Sensorimotor function, evaluated by the modified Garcia test, was impaired in SAH mice up to 1 week with significant deficits observed at 1, 2 and 7 dpi compared to sham (p<0.001) (Fig. 2E). SAH mice also developed significant vasospasm and structural damage. A persistent decrease in the lumen to wall thickness ratio of the middle cerebral artery (MCA) was observed at 3 and 7 dpi (p<0.05 3 dpi and p<0.01 7 dpi, SAH vs sham) (Fig. 1F). Additionally, MRI-based DWI showed increased hyperintense voxels in SAH mice (p<0.01, SAH vs. sham), consistent with vasogenic edema near the site of blood injection (BI) (Fig. 1G).

**Fig. 1.**
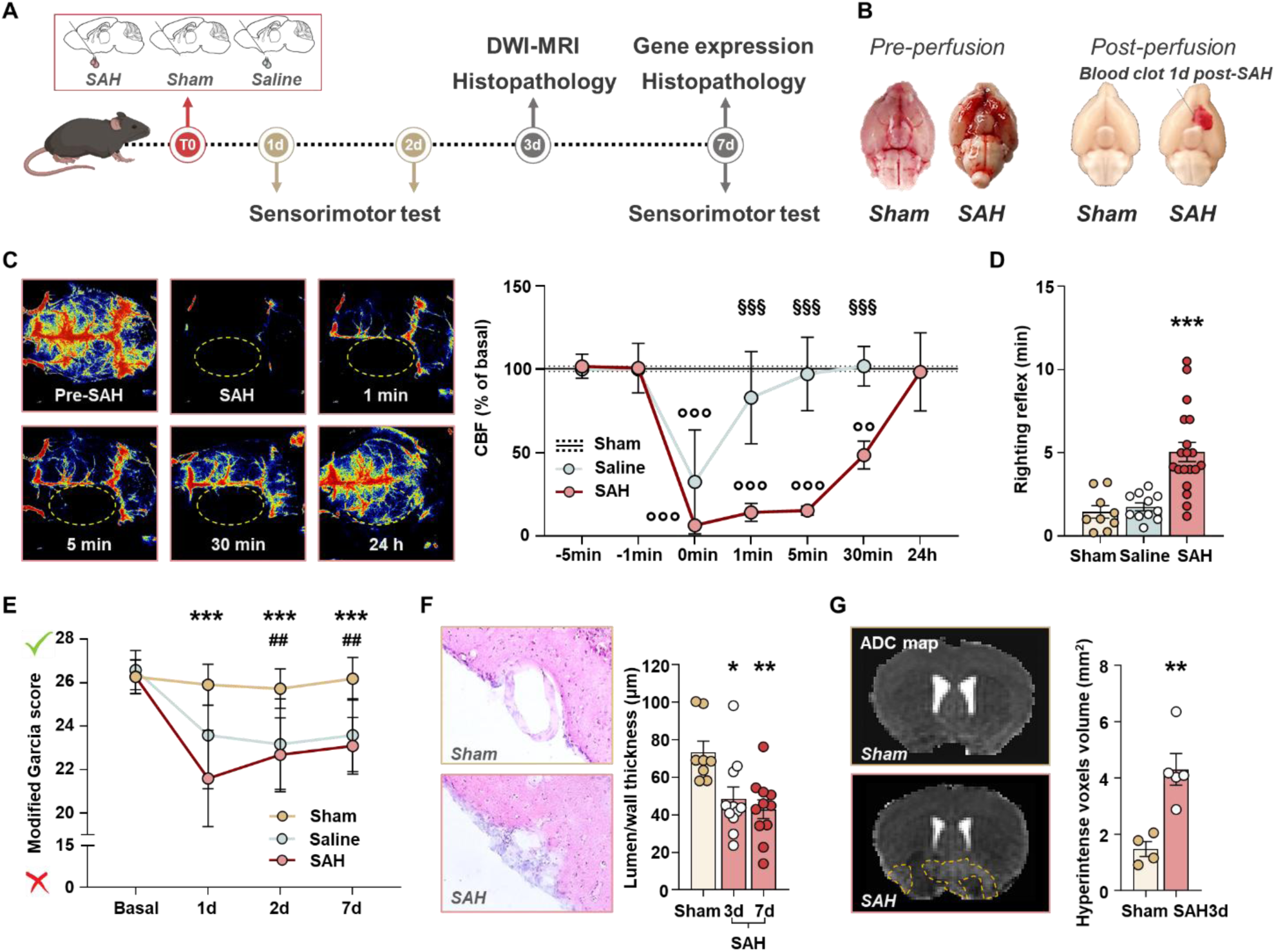
Evaluation of vasospasm, neurological deficits, brain edema in a clinically relevant model of SAH. **A)** Experimental design. SAH (n=36), saline (n=16) and sham (n=24) procedures were performed at T0. **B)** Representative brain images of sham and SAH brains collected immediately after the surgery and post perfusion at 1 dpi. **C)** CBF monitoring by LSCI up to 24 hours following sham (n=3), SAH (n=3) or saline (n=3) injection. Two-way ANOVA, Sidak post hoc test. §§§ p<0.001 SAH vs saline. °°° p<0.001, °° p<0.01 vs basal values. Data are mean ± SD. **D)** Righting reflex latency post-surgery. One-way ANOVA followed by Tukey post hoc test. *** p<0,001 SAH vs sham and SAH vs saline. **E)** Longitudinal evaluation of sensorimotor function by modified Garcia test up to 7 dpi in SAH (n=12) saline (n=12) and sham (n=12) mice. Mixed-effect model followed by Šídák post hoc multiple comparison test. *** p<0.001 SAH vs sham; ## p<0,01, # p<0,05 sham vs saline. Data are mean ± SD. **F)** Representative images and quantification of the lumen thickness ratio of the MCA at 3 and 7 dpi in SAH and sham group. Kruskal-Wallis followed by Dunn’s post hoc test. * p<0.05, ** p<0.01 vs sham. Data are mean ± SEM. **G)** Vasogenic edema by MRI DWI at 3 dpi in SAH and sham group. Dotted yellow line indicates the area of hyperintense voxels. Unpaired T test. ** p<0,01 SAH vs sham. SAH, subarachnoid hemorrhage; LSCI, laser speckle contrast imaging; dpi, days post injury; MCA, middle cerebral artery; MRI, magnetic resonance imaging; DWI, diffusion-weighted imaging; ADC, apparent diffusion coefficient; SD, standard deviation; SEM, standard error of the mean.

**Fig. 2.**
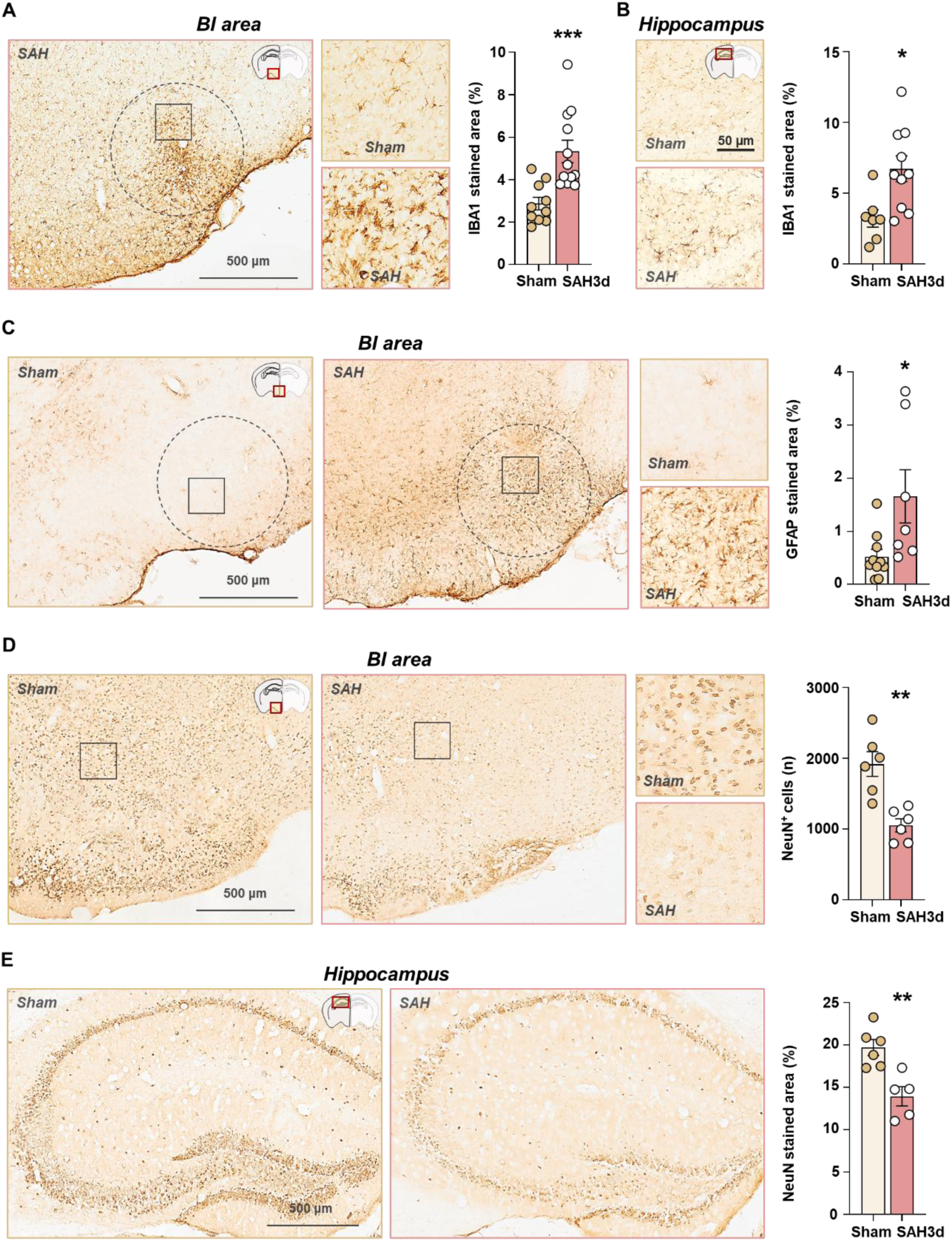
Histological evaluation of acute neuroinflammatory response triggered by SAH. A) Representative images and quantification of IBA1+ stained area in the BI area. Mann Whitney test. *** p<0.001 vs sham. B) Representative images and quantification of IBA1+ stained area in the hippocampus at 3 dpi. Unpaired T test. * p<0.05 vs sham. C) Representative images and quantification of GFAP+ stained area in the BI area at 3 dpi. Mann Whitney test. *** p<0.001 vs sham. D) Representative images and quantification of NeuN+ cell count in the BI area and E) NeuN+ stained area in the hippocampus, at 3 dpi. Unpaired T test. ** p<0,01 vs Sham. Data are mean ± SEM. SAH, subarachnoid hemorrhage; dpi, days post injury; SEM, standard error of the mean; BI area, blood injection area.

### SAH induces acute tissue neuroinflammation characterized by glial reactivity and neuronal death

Gene expression analysis by RT-qPCR revealed upregulation in *Cd11b* (p<0.01) and *Gfap* (p<0.05) genes in the BI area at 3 dpi in SAH mice compared to sham (Fig. S1). Immunohistochemistry further confirmed elevated glial activation. SAH mice exhibited a significantly larger coverage of IBA1+ cells in both the BI area (p<0.05) (Fig. 2A) and the hippocampus (p<0.05) (Fig. 2B), indicating microglial activation. Increased GFAP positivity in the BI area at 3 dpi (p<0.05) also confirmed reactive astrogliosis (Fig. 2C). Neuronal loss accompanied glial reactivity, as shown by a significant reduction in NeuN+ cell at 3 dpi in SAH mice compared to sham both in the BI area (p<0.001) (Fig. 2D) and hippocampus (p<0.01) (Fig. 2E).

### Hippocampal transcriptional changes indicate a major involvement of T cells and microglia in the neuroinflammatory response after SAH

To investigate the neuroinflammatory response to SAH, we performed transcriptomic analysis of the hippocampus, a region distal to the site of blood injection, in SAH, saline and sham mice. We performed a targeted NanoString analysis using a panel encompassing 700 neuroinflammation-related genes. The transcriptomics revealed 22 differentially expressed genes (DEGs) in SAH mice compared to sham and saline controls (−0.5 > log2 ratio > 0.5 and adj p < 0.05) (Fig. 3A, B). Pathway enrichment analysis of these DEGs, based on the proportion of DEGs relative to the total genes within each pathway, identified adaptive immunity, inflammatory signaling and innate immunity as the most prominently regulated pathway (Fig. 3C). Assigning cell type specificity to DEGs revealed that T cells associated genes constituted the largest fraction of the DEGs within the adaptive immunity pathway (∼34%), while microglial genes represented the majority (∼51%) within the innate immunity pathway (Fig. 3D). Importantly, most of DEGs were found in the intersections between the SAH vs sham and SAH vs saline comparisons (Fig. 3E), with few detected in the saline vs sham comparison. This indicates that the majority of the observed transcriptional changes were primarily driven by the presence of blood breakdown products in the subarachnoid space and are specific to SAH pathology. Notably, several microglial DEGs identified in our model (highlighted in orange) have previously been implicated in a disease-associated microglial phenotype that is specifically dependent on T cell–microglia interactions in a murine model of stroke (Benakis *et al*., 2022) (Fig. 3E). In contrast, *Spp1* (highlighted in green), while also a microglial gene, has been shown to be regulated independently of T cell interactions (Benakis *et al*., 2022). *Spp1* was differentially expressed in both SAH vs. sham and saline vs. sham comparisons, suggesting its regulation reflects a general injury response rather than a SAH-specific mechanism. We selected key lymphocytic genes the be validated by qPCR in the hippocampus and to verify their expression in the BI area (Fig. 3F). We confirmed that *Fas* and *Il6ra* were selectively up regulated in the hippocampus of SAH mice. In the BI area the specific T lymphocytic gene *Trac* and *Ccr5* were also upregulated in SAH mice, thus suggesting in increased T cell infiltration in the brain parenchyma close to the BI area.

**Fig. 3.**
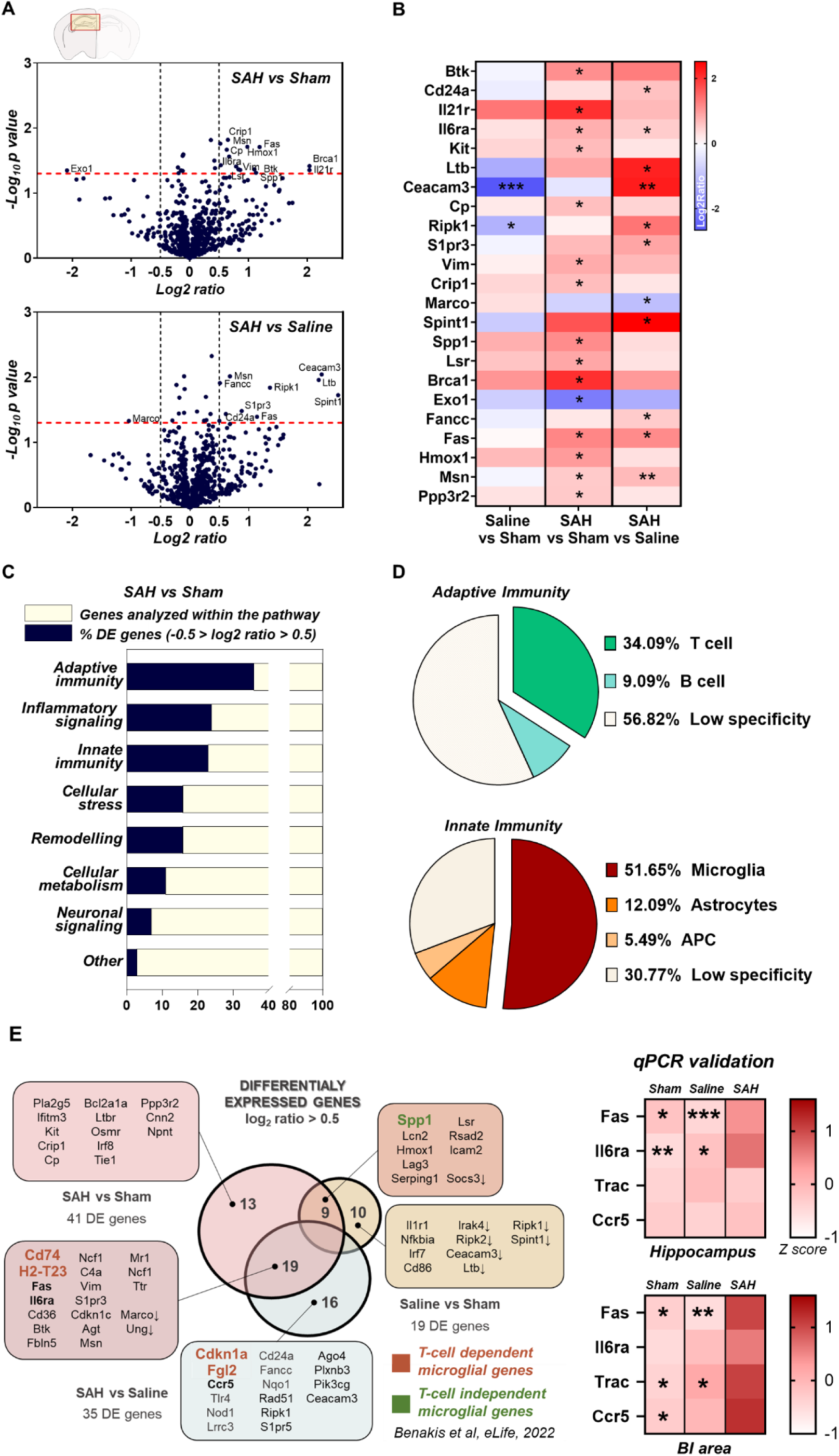
NanoString analysis of neuroinflammatory genes expression in hippocampus at 7 days after SAH. **A)** Volcano plot showing differentially expressed (DE) genes (log10 adj P value > 5, −0.5 > log2 (ratio) > 0.5) in SAH (n=4) vs sham (n=4) (upper plot) and SAH vs saline (n=4) (lower plot). **B)** Heat map showing the log2 ratio of average gene count indicating gene expression changes for significantly DE genes in SAH vs sham and SAH vs saline. Red indicates up-regulated genes and blue indicates down-regulated genes. One-way ANOVA, FDR adjusted p-value. * p<0.05, ** p<0.01, *** p<0.001. **C)** Bar graph showing regulated pathways according to genes found to be DE from the NanoString analysis. **D)** Pie charts illustrating the cell type specificity for the DE genes. **E)** On the left, Venn chart showing DE genes with intersections for each condition. Arrows indicate down regulated genes. On the right, heat map showing qPCR validation for relevant DE genes. Unpaired T test, *** p<0.001, ** p<0.01, * p<0.05 SAH vs sham. SAH, subarachnoid hemorrhage; DE, differentially expressed; qPCR, quantitative polymerase chain reaction, BI area, blood injection area.

### SAH induces T cell recruitment and infiltration into the brain parenchyma

Building on the transcriptional evidence implicating adaptive immune activation, we next examined T cell infiltration at the tissue level using immunofluorescence. CD3^+^ T cells were detected in the brain parenchyma near the BI area in SAH mice at 3 (p<0.05 vs sham) and 7 dpi (p<0.001) but not in sham animals (Fig. 4A). To determine whether these T cells originated from the injected blood, we induced SAH using donor blood from CD3KO mice (p<0.05 vs sham, violet bar in the graph) (Fig. 4A). Even under this condition, infiltrating CD3^+^ T cells were still observed in the parenchyma at 3 dpi. This evidence highlights the active recruitment of T cells from the periphery into the brain parenchyma as a key pathophysiological process occurring as early as 3 dpi. By 7 dpi, the majority of infiltrating T cells expressed CD4 (p<0.001 vs CD4) (Fig. 4B), supporting a dominant role for CD4^+^ T cells during the acute phase of the inflammatory response. Clusters of T cells were also localized to the choroid plexus, a region previously identified as key invasion route for lymphocytes into the brain (Llovera *et al*., 2017; Lazarevic *et al*., 2023), as well as the meninges adjacent to the BI area (Fig. 4C-D).

**Fig. 4.**
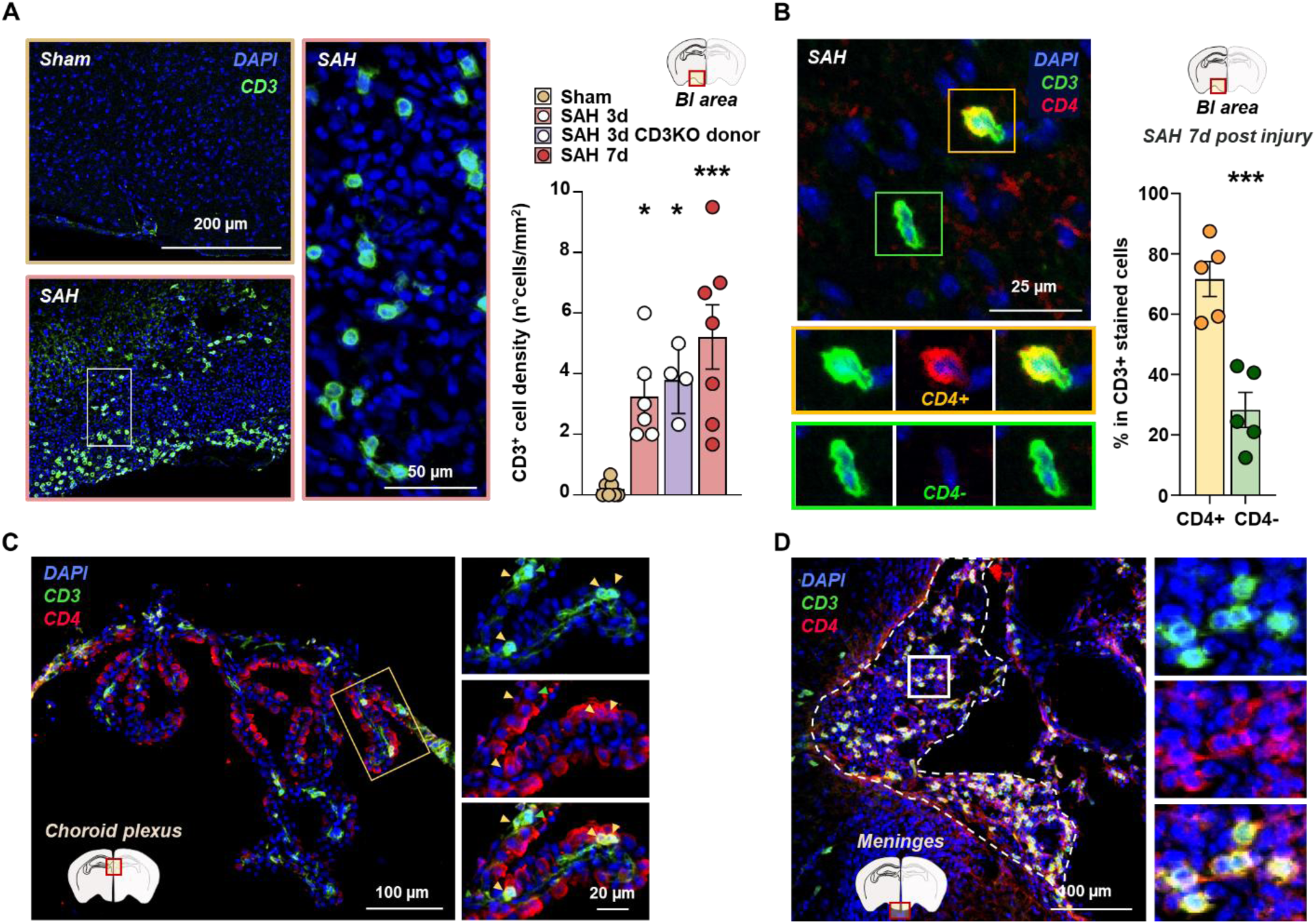
T cell recruitment and infiltration in brain parenchyma after SAH. **A)** Representative images and quantification of CD3+ T cell density in the BI area at 3 and 7 dpi. One-way ANOVA followed by Tukey post hoc test. * p<0,05, ** p<0,01 SAH vs Sham. **B)** Representative images and quantification of CD4+ and CD4-cell rate among CD3+ T cells at 7 dpi. Unpaired T test. *** p<0,001. Data are mean ± SEM. **C)** Representative image of infiltrating T cells in the choroid plexus and **D)** in the meninges at 7 dpi. SAH, subarachnoid hemorrhage; dpi, days post injury; SEM, standard error of the mean, BI area, blood injection area.

### Time-dependent increase of CD4^+^ T cell paralleled by rising levels of NfL and IL-6 in the CNS is a hallmark of SAH in patients

To establish the clinical relevance of T cells involvement observed in the experimental SAH model, we analyzed peripheral blood and CSF by flow cytometry in 46 patients with SAH. Clinical characteristics are summarized in Table 1. We evaluated the temporal dynamics of CD4⁺ and CD8⁺ T cells in the CSF within 48 hours (T1), and between day 5-10 (T2) from SAH. CD4⁺ T cells showed a selective increase in the CSF over time (Fig. 5A), whereas CD8⁺ T cells decreased (Fig. 5B). These findings suggest a predominant role of CD4⁺ T cells in the ongoing brain injury response in SAH patients, consistent with our experimental findings. Notably, CD4⁺, but not CD8⁺ T cells, were also elevated in the peripheral blood of SAH patients compared to healthy donors, indicating a systemic expansion of CD4⁺ T cells (Fig. S2). These temporal changes in CD4+ T cell recruitment were paralleled by rising CSF IL6, but not plasma IL6, pointing to a localized neuroinflammatory process (Fig. 5C). In contrast, NFL increased from T1 to T2 in both blood and CSF, reflecting ongoing neuronal injury (Fig. 5D).

**Fig. 5.**
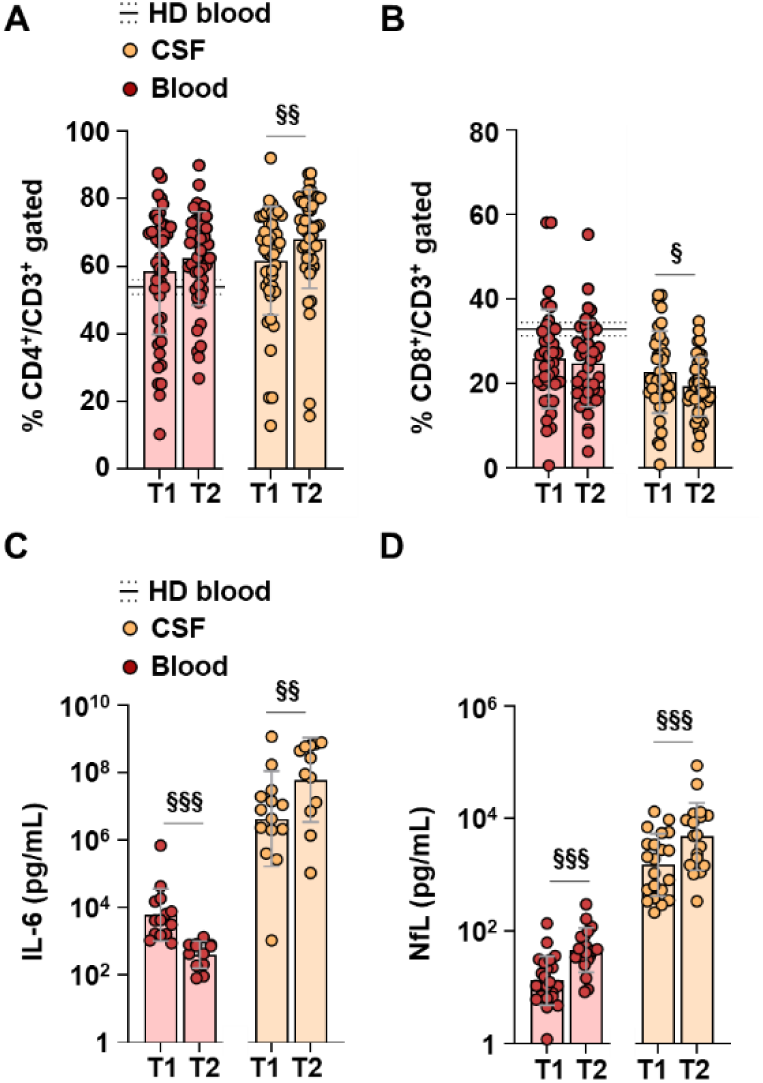
Longitudinal analysis of T cells, NfL and Il6 levels in blood and CSF of SAH patients**. A)** Longitudinal analysis of CD4+ cells and **B)** CD8+ cells frequencies among gated CD3+ cells, in blood (red dots) and CSF (yellow dots) at T1 (48 hours, n=46) and T2 (5-10 days, n=38) in SAH patients. **C**) Longitudinal analysis of IL-6 levels and **D**) NfL in blood (red dots) and CSF (yellow dots) in SAH patients. Wilcoxon matched-pairs signed rank test. § p<0,05, §§ p<0,01, §§§ p<0,001. Data are mean ± SEM. CSF, cerebrospinal fluid; SAH, subarachnoid hemorrhage; dpi, days post injury; IL-6, interleukin 6; NfL, neurofilament light chain; SEM, standard error of the mean.

**Table 1.** Demographic information of SAH patients. WFNS, World federation of neurosurgery society; GOS, Glasgow outcome scale

| Patients characteristic (n=46) |  |
| --- | --- |
| Age (median and range) | 58 (35-84) |
|  | % |
| Gender |  |
| Female | 70 |
| Male | 30 |
| Treatment |  |
| Coiling | 89 |
| Clipping | 11 |
| WFNS |  |
| Severe (4-5) | 65 |
| Mild (1-3) | 35 |
| mFisher |  |
| Grade 1-2 | 11 |
| Grade 3-4 | 89 |
| Intraparenchymal hemorrhage |  |
|  | 30 |
| Radiological vasospasm |  |
|  | 52 |
| Delayed neurological deficit |  |
|  | 30 |
| Delayed cerebral infarction |  |
|  | 28 |
| GOS (6-months) |  |
| Favorable (4-5) | 25 |
| Unfavorable (1-3) | 75 |

### T cell depletion mitigates microglial reactivity and neuronal loss after SAH

To explore the role of T cells in modulating microglial activation and neurodegeneration, we performed gene expression and histopathological analyses in WT and CD3KO SAH mice.

Microglial activation, measured by IBA1 immunoreactivity, was markedly increased in WT SAH mice throughout the BI area, with significant expansion observed up to 1 mm from the injection site (p<0.05 at 0.5 mm, p<0.01 at 1 mm) (Fig. 6A, E). In contrast, CD3KO mice exhibited a more restricted IBA1+ response, confined within 0.5 mm of the core of the BI area (p<0.05), with no significant increase in regions beyond that distance. A morphological transition in microglia between the 0.5 mm and 1 mm zones further highlighted the spatial limitation of the inflammatory response in CD3KO mice (Fig. 6A). In the hippocampus, IBA1+ area coverage was significantly increased in WT SAH mice but not in CD3KO SAH mice (p<0.05) (Fig. 6B, F) indicating a lack of remote microglial activation in the absence of T cells.

**Fig. 6.**
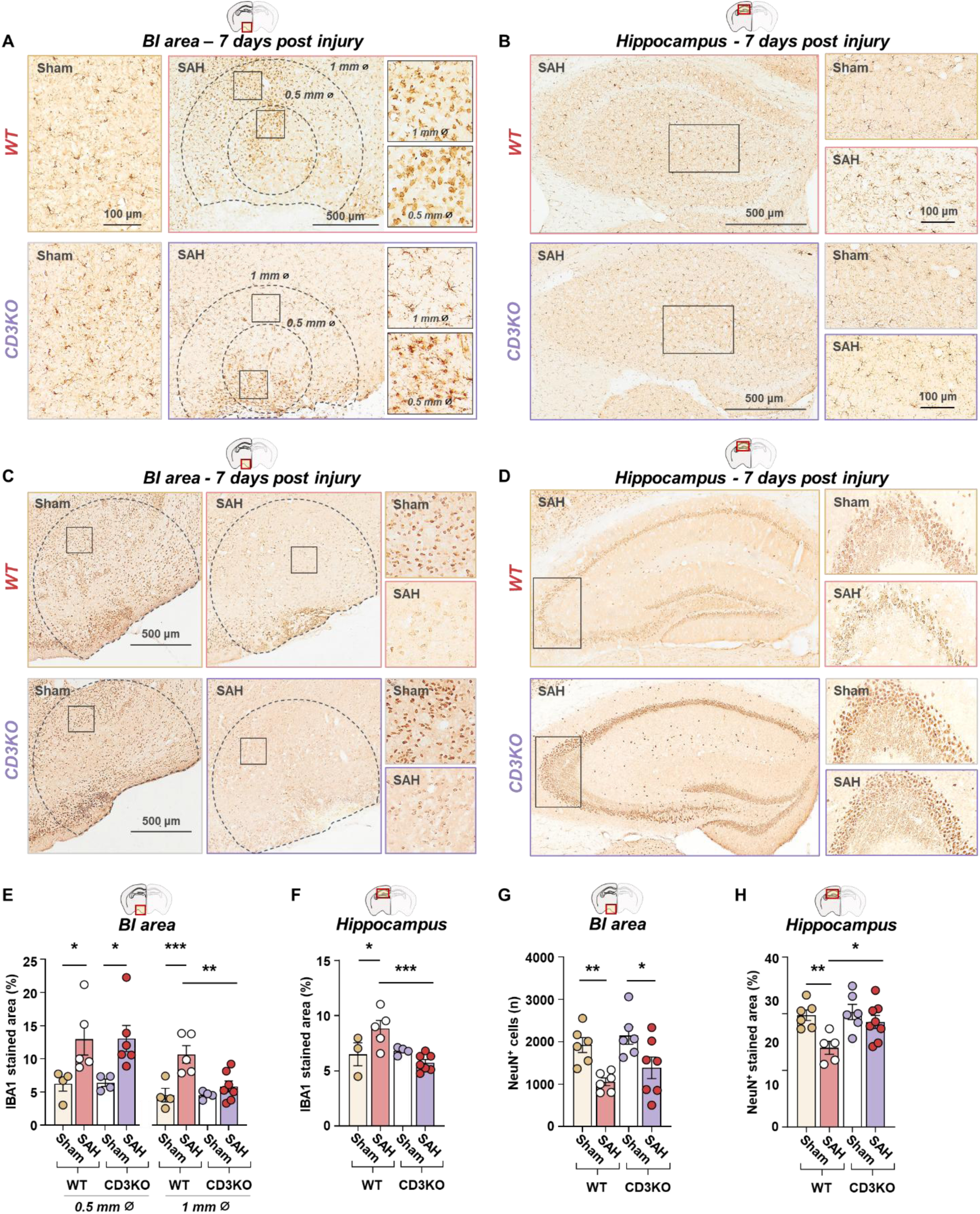
Changes in microglial reactivity and neuronal death in CD3KO mice at 7 dpi. **A)** Representative images of IBA1+ stained area in the BI area and **B)** in the hippocampus, at 3 dpi, in sham or SAH WT and CD3KO mice. **C)** Representative images of NeuN+ cell in the BI area and **D)** NeuN+ stained area in the hippocampus, at 3 dpi, in sham or SAH WT and CD3KO mice. **E)** Quantifications of the percentage of IBA1+ stained area in the BI area and **F)** in the hippocampus, in sham or SAH WT and CD3KO mice. **G)** Quantifications of the NeuN+ cell count in the BI area and F) NeuN+ stained area in the hippocampus, in sham or SAH WT and CD3KO mice. Two-way ANOVA, Fisher’s LSD. *** p<0,001, ** p<0,01, * p<0,05. Data are mean ± SEM. SAH, subarachnoid hemorrhage; IBA1, ionized calcium-binding adapter molecule 1; dpi, days post injury; SEM, standard error of the mean; BI area, blood injection area.

Neuronal loss, quantified by NeuN staining, was evident in both genotypes within the core of the BI area (p<0.01 in WT mice, p<0.05 in CD3KO mice) (Fig. 6C, G). However, significant hippocampal neuronal loss was observed only in WT SAH mice (p<0.01), and not in CD3KO SAH mice (Fig. 6D, H), suggesting that T cells contribute to remote neuronal degeneration. To dissect the molecular basis of this differential glial response, we evaluated expression of microglial genes hypothesized to be either T cell–dependent (*Cd74*, *H2-t23*, *Fgl2*) or T cell–independent (*Spp1*). Confirming T cell-independent regulation, *Spp1* was upregulated in both WT (p<0.01) and CD3KO mice (p<0.06), compared to their respective sham controls. In contrast, *Cd74* (p<0.01), *H2-t23* (p<0.05), and *Fgl2* (p<0.05) were significantly upregulated exclusively in WT SAH mice, with no significant changes observed in CD3KO mice (Fig. 7B, C), demonstrating that their regulation is T cell– dependent. Consistent with the acute reduction of cerebral blood flow induced by SAH, in both CD3KO and WT mice, cerebral vasospasm and neurological deficits were present (Fig. S2). However, CD3KO mice demonstrated a significantly attenuated neuroinflammatory response compared to WT mice following SAH.

**Fig. 7.**
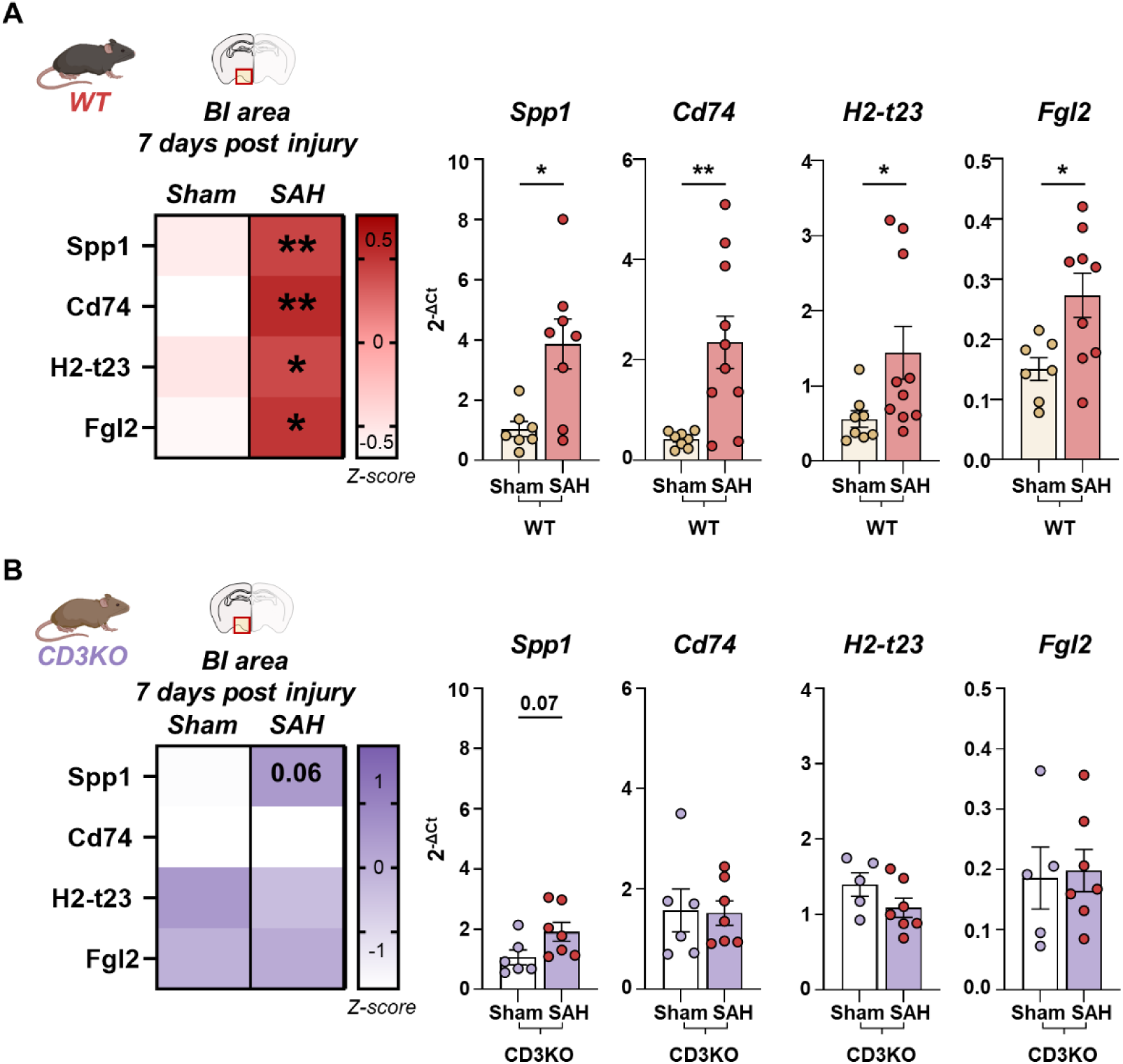
Microglial gene expression profile changing in CD3KO mice after SAH at 7 dpi. **A)** Experimental design. Histopathological analysis was performed at 3 dpi and gene expression analysis was performed at 7 dpi in WT and CD3KO Sham or SAH mice. **B)** Heat map and bar graphs showing qPCR analysis for microglial genes whose expression has been shown to be selectively modulated by T cell interaction, in Sham or SAH WT and **C)** CD3KO mice. Mann Whitney test, ** p<0.01, * p<0.05 SAH vs Sham. Data are mean ± SEM. SAH, subarachnoid hemorrhage; IBA1, ionized calcium-binding adapter molecule 1; GFAP, glial fibrillary acidic protein; dpi, days post injury; SEM, standard error of the mean; BI area, blood injection area.

## Discussion

In this study we evaluated the neuroinflammatory response, focusing on the role of T cells in modulating the secondary injury evolution after SAH. Our translational approach, combining longitudinal analysis in SAH patients with a mechanistically tractable mouse model, revealed that CD4⁺ T cells are actively recruited into the CNS post-SAH, where they contribute to microglial activation and neuronal injury. We demonstrated that SAH patients show a time-dependent increase in CD4⁺ T cells in the CSF, accompanied by rising markers of neuroinflammation (IL-6) and neuronal damage (NfL). Parallel experiments in a clinically relevant murine model recapitulated these human features, validating the model and enabling in-depth mechanistic investigation. Importantly, we show that the neuroinflammatory response in mice is characterized by CD4⁺ T-cell infiltration into the brain parenchyma and that these cells play a key role in shaping the microglial response and exacerbating neuronal loss.

The pre-chiasmatic blood injection SAH mouse model (Sabri *et al*., 2009) offers a high degree of reproducibility and control over the site and extent of hemorrhage, which is critical for studying secondary injury processes. Immediately after SAH, mice exhibited a rapid and prolonged reduction of CBF, consistent with clinical reports of acute ischemia in SAH patients (Carteron *et al*., 2017). Key pathological features (Crowley *et al*., 2011; Macdonald, 2014; Thilak *et al*., 2024), including cerebral vasospasm, vasogenic edema (as detected by DWI-MRI), sensorimotor deficits, and histologically confirmed neuronal death in the parenchymal area close to the BI area, were all evident within the first week post-injury.

Histopathological analysis revealed widespread microglial reactivity and astrogliosis, particularly near the site of blood injection and extending into the hippocampus, a region anatomically remote from the BI area, suggesting a widespread neuroinflammatory cascade. These findings were supported by transcriptomic profiling of the hippocampus, which showed strong enrichment of both innate and adaptive immune pathways, with T cell–associated genes dominating the adaptive response and microglial genes dominating the innate response. Importantly, using the flexibility of our model to inject either blood or saline into the subarachnoid space, we demonstrated that this inflammatory response was specifically driven by the presence of blood. Only SAH mice exhibited upregulation of microglial genes associated with a disease-associated phenotype including those shown to be modulated by T cells (Benakis *et al*., 2022). This supports a model in which blood breakdown products initiate a T cell–dependent amplification loop, exacerbating microglial activation and neuronal damage. While it is well know that pro-inflammatory T cell subsets such as Th1 and Th17 can promote neurotoxicity (Shichita *et al*., 2009; González and Pacheco, 2014) whereas regulatory T cells can suppress these population, and exert neuroprotective effects (Yshii *et al*., 2022). Evidence from ischemic stroke model shows that complete T cell depletion reduces lesion size and improves neurological function (Kleinschnitz *et al*., 2010). Moreover, passive immune CD3 suppression has been shown to produce neuroprotective effects after experimental TBI, suggesting a net deleterious effect of T cell–mediated inflammation during acute CNS injury (Izzy *et al*., 2025).Our histopathological data confirmed the presence of CD3⁺ T cells infiltrating the brain parenchyma post-SAH, with a predominance of CD4⁺ cells by 7 dpi. Importantly, using donor blood from CD3 knockout (CD3KO) mice to induce SAH, we showed that these T cells did not originate from the injected blood, but were instead recruited from the host immune system. This active recruitment reinforces the role of T cells as integral effectors of the post-SAH immune response.

Patient CSF analysis supported these findings, showing a selective increase in CD4^+^, but not CD8^+^, T cell levels from 1 to 7 dpi. Few observational studies show CD4^+^ T cells displaying dynamic changes in peripheral blood and in the CSF of SAH patients. Interestingly, while prior studies in other acute brain injuries (e.g., stroke, TBI) have reported systemic alterations in both CD4⁺ and CD8⁺ compartments (Xie *et al*., 2019; Magatti *et al*., 2023), our data show that SAH selectively drives expansion of CD4⁺ T cells. This suggests disease-specific immune dynamics and highlights the potential role of CD4^+^ T cell in shaping injury evolution after SAH. In particular, the imbalance between pro-inflammatory CD4^+^ IL-17A secreting T cells is reported to be associated with the development of vasospasm in SAH patients (Moraes *et al*., 2020; Romoli *et al*., 2023) though these findings require validation in larger cohorts. The lack of access to brain tissue in patients underscores the value of our animal model in dissecting immunopathological mechanisms at the tissue level. To determine the functional role of T cells in modulating the SAH response (Xu *et al*., 2015; Dong *et al*., 2021; Li *et al*., 2025), we compared wild-type and CD3KO mice. Although both groups exhibited similar neurological function and vasospasm, CD3KO mice showed significantly reduced microglial activation and neuronal loss. This was particularly evident in the hippocampus, where wild-type mice displayed extensive pathology that was absent in CD3KO miceGene expression analysis further clarified the immunological interplay. We found that *Spp1*, a microglial gene known to be regulated independently of T cells, was upregulated in both wild-type and CD3KO mice. In contrast, *Cd74*, *H2-t23*, and *Fgl2*, genes previously implicated in T cell microglia crosstalk (Benakis *et al*., 2022), were selectively upregulated in wild-type mice but remained unchanged in CD3KO mice. Importantly, recent studies have shown that targeting microglial pro- inflammatory activation could help reduce neuronal damage after SAH (Mao *et al*., 2024; Sun *et al*., 2024; Cui *et al*., 2025). Although CD8^+^ T cells have also been implicated in the pathophysiology of SAH, primarily in promoting BBB disruption following SAH (Li *et al*., 2025), our data support a dominant role for CD4⁺ T cells in modulating microglial responses. This aligns with reports in ischemic models where CD4⁺ T cells drive microglial polarization toward a proinflammatory phenotype (Benakis *et al*., 2022; Shi *et al*., 2023).

Given the deleterious effects of T cell infiltration, therapeutic strategies aimed at modulating the adaptive immune response, such as selective inhibition of proinflammatory T cells or modulation of their inflammatory phenotype, could provide a novel approach to mitigating SAH-induced brain injury. Future studies should explore the molecular mechanisms through which CD4^+^ T cells interact with microglia and evaluate immunomodulatory therapies in SAH preclinical models.

## Conclusion

Our study provides compelling evidence that T cells, particularly the CD4^+^ T helper subset, are key drivers of neuroinflammation and secondary neuronal injury following SAH. The attenuation of inflammatory responses and neuronal loss in CD3KO mice directly implicates T cells as pathogenic contributors. These findings not only reinforce the importance of neuroinflammation as a target for therapeutic intervention but also lay the groundwork for future strategies aimed at modulating adaptive immune responses to improve outcomes in SAH patients.

## Materials and methods

### Mice

To comprehensively characterize the temporal and mechanistic features of neuroinflammation following SAH, a total of 60 adult (9 weeks) male mice were randomly allocated across two experimental cohorts. The first cohort (n=36) was designed to evaluate pathophysiological outcomes up to 7 days post injury (dpi), including vasospasm, edema, sensorimotor deficits, and immune cell infiltration. Animals were assigned to three groups: sham (n=12), saline control (n=12), and SAH (n=12). The second cohort (n=24) was established to assess the specific contribution of T cells to post SAH neuroinflammation and neuronal damage, using CD3 knockout (CD3KO) mice. This cohort included sham WT (n=6), SAH WT (n=6), sham CD3KO (n=6), and SAH CD3KO (n=6) groups. C57BL/6J mice (Envigo, Italy) and CD3KO mice (B6;129-Cd3etm1Lov/J;Jackson Laboratory, USA) were housed in a specific pathogen-free vivarium at a constant temperature (21 ± 1 °C) and relative humidity (60 ± 5%) with a 12h light/dark cycle and free access to pellet food and water. All animal experiments were designed in accordance with the ARRIVE guidelines, with a commitment to refinement, reduction, and replacement, minimizing the numbers of mice, and using biostatistics to optimize mouse numbers on the bases of our preliminary result. Procedures involving animals and their care were conducted in conformity with the institutional guidelines at the IRCCS – Mario Negri Institute for Pharmacological Research in compliance with national (D.lgs 26/2014; Authorization n. 19/2008-A issued March 6, 2008 by Ministry of Health) and international laws and policies (EEC Council Directive 2010/63/UE; the NIH Guide for the Care and Use of Laboratory Animals, 2011 edition). They were reviewed and approved by the Mario Negri Institute Animal Care and Use Committee that includes ad hoc members for ethical issues and by the Italian Ministry of Health (Decreto no. 141/2021-PR). Animal facilities meet international standards and are regularly checked by a certified veterinarian who is responsible for health monitoring, animal welfare supervision, experimental protocols and review of procedures. Behavioral, imaging, and histological assessments were performed by researchers blinded to the experimental groups.

### Experimental subarachnoid hemorrhage

The SAH was performed by a single injection of blood into the pre-chiasmatic cistern, as previously described (Sabri *et al*., 2009). Mice were anesthetized by inhalation of isoflurane (induction 3%; maintenance 1.5%) in a mixture of N_2_O/O_2_ (70%/30%) and placed in a stereotactic apparatus. Corneas were protected by applying lubricant gel during surgery. The skin was disinfected and sagittal incised with a scalpel for a length of 1-2 cm at the midline of the skull. During all surgical procedures, mice were maintained at a body temperature of 37°C. A hole was drilled in the skull 4.5 mm anterior to the bregma. A 27-G spinal needle was advanced at a 24° angle through the hole to the base of the skull. Once the base of the skull was contacted, the needle was retracted by 0.5 mm to proceed with the injection of 80 μl of whole non-heparinized homologous blood (SAH group) or saline (saline group) in the pre-chiasmatic cistern. Sham-operated mice received identical anesthesia and surgery without any injection. All the animals were placed in a recovery cage after surgery in order to monitor their awakening.

### Behavioral test

#### Modified Garcia Test for sensorimotor function

Sensorimotor deficits were assessed in a blinded manner after SAH using the Garcia test (Garcia, Liu and Ho, 1995), modified to include the evaluation of hind paw outstretching, interaction and head tilt. A score from 3 (normal) to 0 (severely deficient) was assigned for each of the following indices: 1) spontaneous activity; 2) response to vibrissae stimulation; 3) response to flank stimulation; 4) body proprioception; 5) climbing; 6) forepaw outstretching; 7) hind paw outstretching; 8) interaction; 9) head tilt; The maximum score achievable by each animal was 27.

### Laser Speckle Contrast Imaging

LSCI was carried out using a RFLSI laser speckle (RWD Life Science, China) according to manufacturer’s instructions as described in a previous study (Deng *et al*., 2025). Speckle patterns were generated and recorded at a frame rate of 15 images per sec. LSCI baseline images were obtained for a period of 5 min before SAH and continued for 30 min and again at 24h. All measurements were performed under anesthesia with a mixture of N_2_O/O_2_ (70%/30%) and placed in a stereotactic apparatus while maintaining the body temperature at 37 °C on a constant heating pad. A midline incision was made to expose the calvaria. The corresponding images were analyzed using the software installed in the laser speckle imaging system. For data analysis, images were averaged over 1 minute for a single region of interest (ROI) covering the left hemisphere. This allowed us to calculate the % change in cerebral perfusion following compared to baseline.

### MRI acquisition and analysis

Anaesthetized mice (isoflurane 1.5 - 2 vol% in an N_2_O/O_2_ (70%/ 30%) were positioned in the magnet. Respiratory frequency was monitored throughout the experiment and body temperature was maintained at 37°C with a heating pad. Brain imaging was done on a 7T small-bore animal scanner (BioSpec®; Bruker, Ettlingen, Germany) running ParaVision 6.01 and equipped with a quadrature ^1^H CryoProbe™ (Bruker, Ettlingen, Germany) surface coil as transmitter and receiver. Diffusion-weighted imaging (DWI), were obtained to quantify edema as previously described (Moro *et al*., 2021). Echo-planar images (EPI) were acquired (TR/TE 2000/48.5 ms, resolution 125 × 125 μm2; FOV 1. 5 × 1.5 cm2; acquisition matrix 120 × 120, slice thickness 0.3 mm) to obtain apparent diffusion coefficient (ADC) maps. Diffusion-encoding was applied in three orthogonal directions with b values of 200, 600 and 900 ms, respectively. ADC-maps were calculated on a pixel-by-pixel basis with an in-house MATLAB script using the model function: ln(S(b)/S0)=-b⋅ADC, where S(b) is the measured signal intensity at a specific b value (b) and S0 the signal intensity in the absence of a diffusion gradient (b = 0). The hyperintense region was traced manually by an expert operator blind to the experimental condition.

### Biochemical analysis

#### NanoString analysis

Mice were euthanized by deep anesthesia (ketamine 150 mg kg-1 and medetomidine 2 mg kg-1, i.p.) and perfused transcardially with 30 ml of phosphate-buffered saline (PBS) 0.1 mol L^-1^. After perfusion the brains were rapidly removed. Blood injection (BI) area, hippocampus and cortex were dissected, immediately frozen in dry ice and stored at −80°C until analysis. RNA was extracted and purified from frozen hippocampal murine tissue using TRI Reagent RT (MRC) and Direct-zol RNA microprep kit (Zymo Research) according to the manufacturer’s instructions. 50 ng of total RNA was probed using the nCounter Mouse Neuroinflammation panel (NanoString Technologies, Seattle, WA), profiling 770 genes across six fundamental themes of neurodegeneration: neurotransmission, neuron-glia interaction, neuroplasticity, cell structure integrity, neuroinflammation, and metabolism. Pathway and cell-type specificity analyses were performed using the pathway and cell-type annotations included in the nCounter Mouse Neuroinflammation panel. For each pathway, the percentage of DE genes was calculated relative to the total number of genes analyzed within that pathway.

#### RT-qPCR

cDNA was synthesized from total RNA by reverse transcription using the qScript cDNA SuperMix (Quanta Bioscience). Target genes were amplified using PerfeCTa SYBR Green FastMix (Quanta Bioscience) and QuantStudio 3 Real-Time PCR System (Thermo Fisher Scientific). Data were normalized to the expression of the *Tbp* housekeeping gene and analyzed using the 2^−ΔCt^ method.

### Histology

Mice were euthanized by deep anesthesia (ketamine 150 mg kg^-1^ and medetomidine 2 mg kg^-1^, i.p.) and perfused transcardially with 30 ml of PBS 0.1 mol L^-1^, pH7.4, followed by 60 ml of chilled paraformaldehyde 4% in PBS. Brains were removed and post fixed for 2 h in 4% PAF, transferred to 20% sucrose in PBS, then frozen in n-pentane for 3 min at −45°C, and stored at − 80 °C until assay. Frozen brains were firstly cut in 20 µm sagittal sections (6 sections in a range between +3,5 mm and +3 mm from midline), then turned and cut in 20 µm coronal sections (10 sections in a range between +1 mm and +0,6 mm from bregma) on a cryostat. Sagittal sections were stained with hematoxylin and eosin for the evaluation of the vasospasm of the middle cerebral artery. Coronal sections were immuno-stained with the following primary antibodies: anti-CD3 (1:500; Rat anti-mouse; BD Pharmigen), anti-CD4 (1:200, polyclonal rabbit anti-mouse, Invitrogen), anti-IBA1 (1:200; Rabbit anti-mouse; DBA Italia), anti-GFAP (1:2000; Mouse anti-mouse; Chemicon), anti-NeuN (1:500, Mouse anti-mouse; Chemicon). The immunoreactions were visualized by indirect detection using the following secondary antibodies: Biotin-goat anti-rabbit (1:200, Vector), Biotin-goat anti mouse (1:200, Vector), Biotin-goat anti-rat (1:200, Jackson Immuno research) using 3,3’-diaminobenzidine (Dako) as chromogen for immunohistochemistry, TSA biotin system (Akoya bioscence) for signal amplification with fluorotyramide as chromogen; Alexa Fluor anti-rat 488 and Alexa Fluor anti-rabbit 647 (all 1:500; Invitrogen) for immunofluorescent visualization without amplification. Nuclei were stained using 4’,6-Diamidin-2-phenylindol (DAPI, Abcam, ab285390) diluted 1:5000 in PBS for 15 minutes at room temperature.

### IHC and immunofluorescence images acquisition and quantification

Bright field images were acquired at 20x with Olympus BX-61-VS microscope, interfaced with VS-ASW-FL software (Olympus, Tokyo); immunofluorescent images were acquired at 40x with Nikon A1 confocal scan unit managed by NIS-Elements software (Nikon, Tokyo, Japan) with 10% overlapping for stitching, z-axis 10 mm. All the images were analyzed using Fiji software (https://fiji.sc/).

For vasospasm quantification, analysis was performed on medio-lateral sections by measuring the vessel lumen area and the average wall thickness of the middle cerebral artery, and expressing the data as a lumen-to-thickness ratio (Sabri *et al*., 2009). BI area was defined in the area of parenchyma proximal to the injection of blood in the pre-chiasmatic cistern. A circular ROI was drawn with a diameter of 1 mm. The center of the circular ROI was placed 0.5 mm below the anterior commissure, based on the injection coordinates. IBA1 stained area was quantified either across the entire BI area or, in the second cohort, within two distinct ROIs: the full BI area (1 mm diameter) and the core of BI area (concentric 0.5 mm diameter), to compare SAH WT and SAH CD3KO mice. GFAP stained area was quantified in the BI area. Neuronal viability was evaluated by performing NeuN^+^ cell count in the BI area and by quantifying NeuN stained area in the molecular layer of hippocampus. CD3^+^ and CD3^+^CD4^+^ cell density was manually calculated in the BI area.

### Human sample collection and analysis

Human subjects were recruited at Fondazione IRCCS Ca’Granda - Ospedale Maggiore Policlinico, in Milan (Italy). Inclusion criteria were aneurysmal subarachnoid hemorrhage diagnosis, hospital admission within 24 hours from the bleeding, age > 18 years and placement of an external ventricular or lumbar drainage for hydrocephalus or intracranial pressure management. Exclusion criteria included pregnancy, previous severe neurological disability and immunological disease. The study was approved by the local ethics committee (Comitato Etico Milano Area 2, Authorization n. 87_2021). All healthy donors were recruited at the Ospedale Maggiore Policlinico, in Milan (Italy) with the approval of the local ethics committee (Comitato Etico Milano Area 2, Authorization n. 708_2020). The treatment protocol for these patients did not differ from standard clinical practice and included early treatment (within 24 hours of admission) of the cerebral aneurysm either by neurosurgical intervention (clipping) or endovascular procedure (coiling), along with evacuation of intraparenchymal hematomas if clinically indicated. Additionally, patients underwent placement of an external ventricular drain (EVD) or lumbar drainage in the presence of clinical or radiological signs of hydrocephalus and/or for intracranial pressure (ICP) monitoring. Simultaneous blood and cerebrospinal fluid (CSF) samples were obtained at two different time-points: within 48 hours from SAH (T1), and between day 5-10 from SAH (T2). Sixty-three consecutive patients were enrolled after admission to the Neuro Intensive Care Unit. However, a total of 17 patients were excluded from the cytometry analysis: in six patients the cellularity of the CSF was inadequate; in the other eleven patients it was impossible to collect the T2 sample (the patients died or the EVD was removed before day 5). Blood samples were collected in EDTA tubes. CSF was obtained from the EVD using sterile technique. 15-20 mL of CSF were collected and transferred in a sterile tube for cytometric analysis. Additionally, three milliliters of CSF and blood samples were centrifuged (2500 rpm for 10 minutes), and from the supernatant, 5 aliquots of CSF (each 500 μL) and 5 aliquots of plasma (each 500 μL) were prepared and stored at −80°C as previously described. These aliquots were used for Neurofilament Light Chain (NfL) and Interleukin −6 analysis.

T cells were isolated from blood and CSF using a Ficoll-Paque density gradient and subsequently stained to identify lymphocyte subpopulations. Each sample was stained with the following markers: anti-CD3 (1:135, mous anti-human, Becton Dickinson), anti-CD4 (1:180, mouse anti-human, Becton Dickinson), anti-CD8 (1:200 mouse anti-human, Becton Dickinson). Data obtained from cytometry were imported into FlowJo software to compensate for fluorescence spreading, and dead cells and debris were excluded from the analyses. T cells were identified as CD3+ cells, forming the basis for the subsequent analysis of all other subpopulations.

Il6 was analyzed by Luminex assay (kit LXSAHM, Luminex® Discovery Assay, R&D System, USA) on Luminex FLEXMAP 3D analyzer (R&D System, USA); NfL levels were analyzed using commercially available single molecule array assay kits (103400, Quanterix, Billerica, MA) on an SR-X Analyzer (Quanterix, Billerica, MA). We followed the manufacturer’s directions, samples were run in duplicate, and experiments were conducted by researchers blinded to the experimental conditions.

### Statistical analysis

GraphPad Prism 10 was used for statistical analyses. Assumptions of normality were checked using the Shapiro-Wilk omnibus test. Outliers were identified using the ROUT method and excluded from the analysis. Two-way ANOVA for repeated measures was used in the case of longitudinal assessments in mice (behavioral tests and CBF), or by one-way ANOVA in the case of three group comparison followed by Tukey’s multiple comparison test. For a two group comparison data were analyzed by an unpaired t test. When the assumption of normality was not met, correspondent non-parametric tests were applied. For each experiment, the figure legend reports the statistical analysis of data. P-values of 0.05 were considered statistically significant. Graphs show the mean + standard error of the mean (SEM) or median and interquartile range. Targeted transcriptomic analysis of the mouse neuroinflammation panel was performed using the nSolver analysis software (v4.0; NanoString Technologies, Inc.), with a threshold cut-off set at >20 normalized average counts. Genes with a log_2_ fold change >0.5 or <−0.5 were considered differentially expressed (DE).

### Data sharing

All data from this manuscript will be publicly available through the open science services Zenodo

## Acknowledgements

This work was supported by the Italian Ministry of Health (Young Investigators Award 2019, GR-2019-12369998), and the Italian Cariplo Foundation (Biomedical Research conducted by Young Researchers, 2019-1632). The authors would like to thank the Fondazione Mario Negri Swiss for supporting this work by fostering collaboration between the Mario Negri Institute and the IRB (Università della Svizzera italiana), and for funding the SR-X™ Biomarker Detection System (Simoa® Technology, Quanterix) used for biomarker analysis in SAH patients. The Department of Pathophysiology and Transplantation, University of Milan, is funded by the Italian Ministry of Education and Research (MUR): Dipartimenti di Eccellenza Program 2023 to 2027.

## Supplementary material

**Fig. S1.**
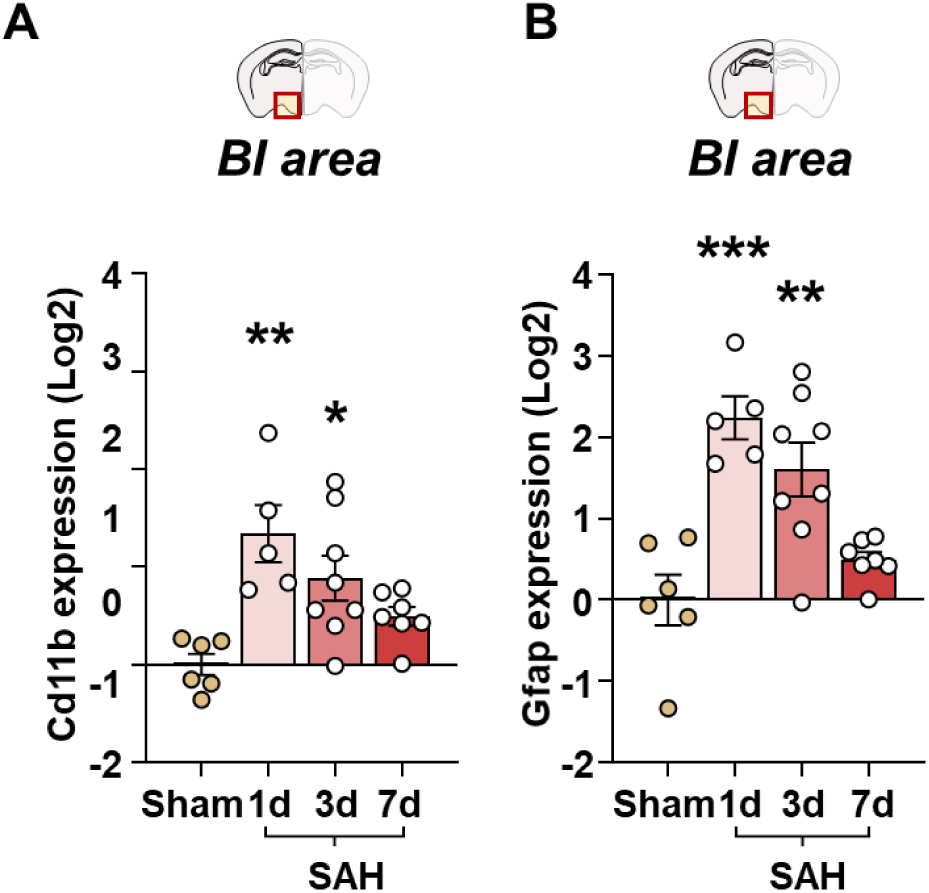
Expression levels of glial genes Cd11b and Gfap in the BI area. **A)** Bar graph showing gene expression level of the glial markers Cd11b and **B)** Gfap evaluated by RT-qPCR in the BI area of Sham and SAH mice 1 and 3 dpi. One-way ANOVA followed by Tukey post hoc test. * p<0,05; ** p<0,01; p<0,001 vs Sham. Data are mean ± SEM. SAH, subarachnoid hemorrhage; dpi, days post injury; SEM, standard error of the mean.

**Fig. S2.**
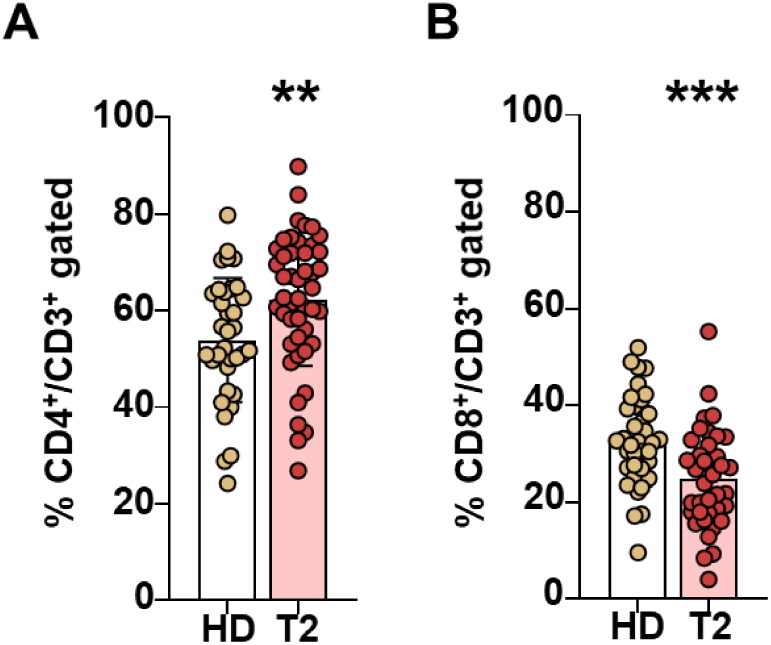
Flow cytometry analysis of T cells levels in the peripheral blood after SAH. **A)** Quantification of CD4+ cells and **B)** CD8+ cells frequencies among gated CD3+ cells, in blood (red dots) and healthy donor (HD, gold dots) at T2 (5-10 days). Unpaired T-test. ** p<0,01, *** p<0,001. Data are mean ± SEM. SAH, subarachnoid hemorrhage; SEM, standard error of the mean.

**Fig. S3.**
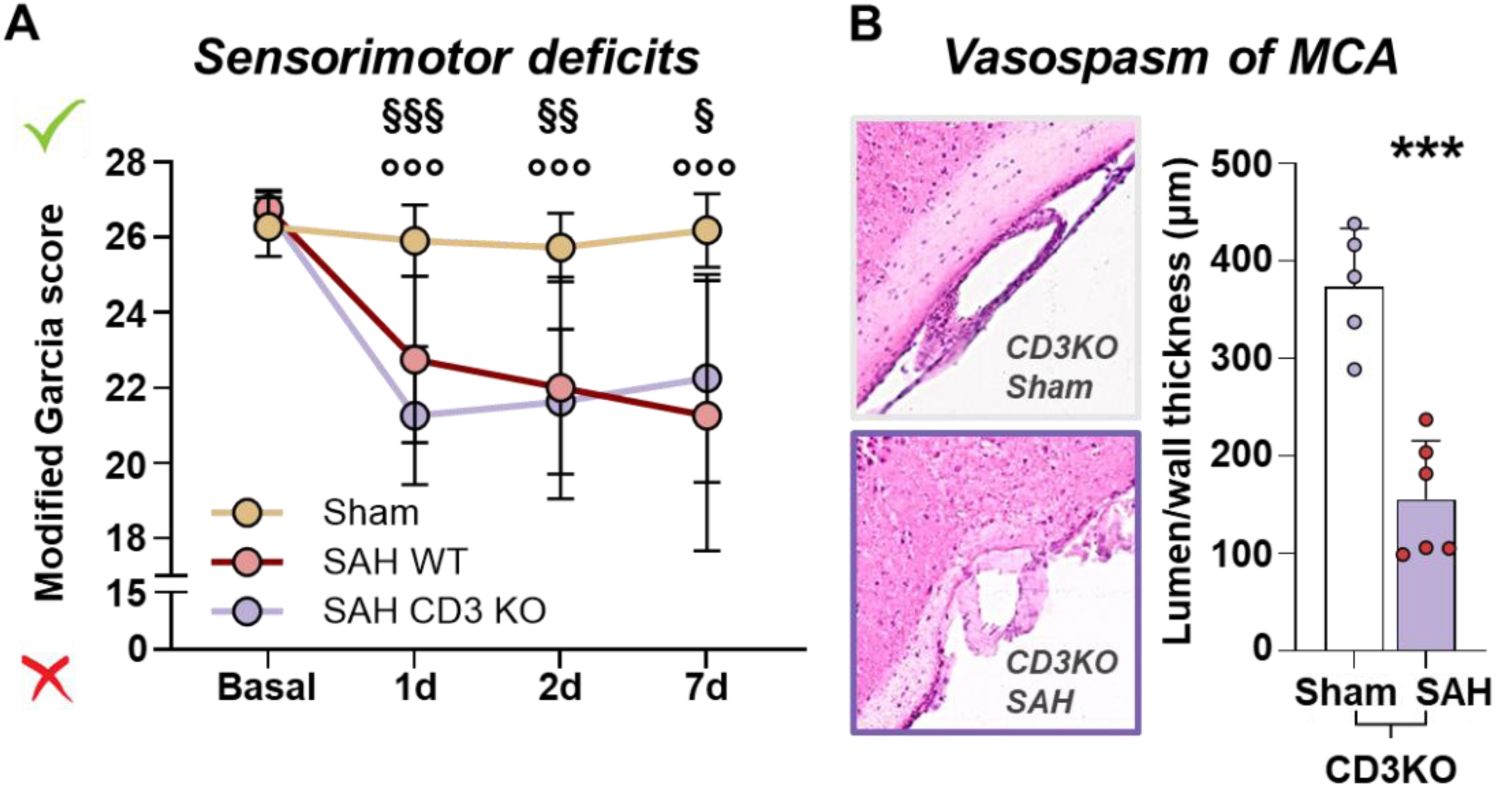
Sensorimotor deficits and vasospasm of middle cerebral artery in CD3KO mice. A) Longitudinal evaluation of the sensorimotor function by Modified Garcia Test up to 7 dpi. Mixed-effect model followed by Šídák post hoc multiple comparison test. §§§ p<0,001, §§ p<0,01, § p<0,05 SAH CD3KO vs Sham; °°° p<0,001 SAH WT vs Sham. Data are mean ± SD. B) Evaluation of vasospasm of the MCA at 7 dpi in SAH CD3KO and Sham group. One-way ANOVA followed by Tukey post hoc test. *** p<0.001 vs Sham. Data are mean ± SEM. SAH, subarachnoid hemorrhage; dpi, days post injury; SD, standard deviation; SEM, standard error of the mean.

## References

Al-Khindi, T., Macdonald, R.L. and Schweizer, T.A. (2010) “Cognitive and Functional Outcome After Aneurysmal Subarachnoid Hemorrhage,” Stroke, 41(8). Available at: 10.1161/STROKEAHA.110.581975.

Benakis, C. et al. (2022) “T cells modulate the microglial response to brain ischemia,” eLife, 11, p. e82031. Available at: 10.7554/eLife.82031.

Carteron, L. et al. (2017) “Non-Ischemic Cerebral Energy Dysfunction at the Early Brain Injury Phase following Aneurysmal Subarachnoid Hemorrhage,” Frontiers in Neurology, 8, p. 325. Available at: 10.3389/fneur.2017.00325.

Chaudhry, S.R. et al. (2021) “Differential polarization and activation dynamics of systemic T helper cell subsets after aneurysmal subarachnoid hemorrhage (SAH) and during post-SAH complications,” Scientific Reports, 11(1), p. 14226. Available at: 10.1038/s41598-021-92873-x.

Chen, S. et al. (2014) “Controversies and evolving new mechanisms in subarachnoid hemorrhage,” Progress in Neurobiology, 115, pp. 64–91. Available at: 10.1016/j.pneurobio.2013.09.002.

Clatterbuck, R.E. et al. (2003) “Prevention of cerebral vasospasm by a humanized anti-CD11/CD18 monoclonal antibody administered after experimental subarachnoid hemorrhage in nonhuman primates,” Journal of Neurosurgery, 99(2), pp. 376–382. Available at: 10.3171/jns.2003.99.2.0376.

Crowley, R.W. et al. (2011) “Angiographic Vasospasm Is Strongly Correlated With Cerebral Infarction After Subarachnoid Hemorrhage,” Stroke, 42(4), pp. 919–923. Available at: 10.1161/STROKEAHA.110.597005.

Cui, Y. et al. (2025) “Silybin attenuates microglia-mediated neuroinflammation via inhibition of STING in experimental subarachnoid hemorrhage,” International Immunopharmacology, 151, p. 114305. Available at: 10.1016/j.intimp.2025.114305.

De Oliveira Manoel, A.L., et al. (2016) “Functional Outcome After Poor-Grade Subarachnoid Hemorrhage: A Single-Center Study and Systematic Literature Review,” Neurocritical Care, 25(3), pp. 338–350. Available at: 10.1007/s12028-016-0305-3.

De Oliveira Manoel, A.L. and Macdonald, R.L. (2018) “Neuroinflammation as a Target for Intervention in Subarachnoid Hemorrhage,” Frontiers in Neurology, 9, p. 292. Available at: 10.3389/fneur.2018.00292.

Deng, H.-J. et al. (2025) “The sentinel against brain injury post-subarachnoid hemorrhage: efferocytosis of erythrocytes by leptomeningeal lymphatic endothelial cells,” Theranostics, 15(6), pp. 2487–2509. Available at: 10.7150/thno.103701.

Di Sapia, R. et al. (2021) “In-depth characterization of a mouse model of post-traumatic epilepsy for biomarker and drug discovery,” Acta Neuropathologica Communications, 9(1), p. 76. Available at: 10.1186/s40478-021-01165-y.

Dong, G. et al. (2021) “Low-Dose IL-2 Treatment Affords Protection against Subarachnoid Hemorrhage Injury by Expanding Peripheral Regulatory T Cells,” ACS Chemical Neuroscience, 12(3), pp. 430–440. Available at: 10.1021/acschemneuro.0c00611.

Fee, D. et al. (2003) “Activated/effector CD4+ T cells exacerbate acute damage in the central nervous system following traumatic injury,” Journal of Neuroimmunology, 136(1–2), pp. 54–66. Available at: 10.1016/S0165-5728(03)00008-0.

Garcia, J.H., Liu, K.-F. and Ho, K.-L. (1995) “Neuronal Necrosis After Middle Cerebral Artery Occlusion in Wistar Rats Progresses at Different Time Intervals in the Caudoputamen and the Cortex,” Stroke, 26(4), pp. 636–643. Available at: 10.1161/01.STR.26.4.636.

Gate, D. et al. (2021) “CD4^+^ T cells contribute to neurodegeneration in Lewy body dementia,” Science, 374(6569), pp. 868–874. Available at: 10.1126/science.abf7266.

González, H. and Pacheco, R. (2014) “T-cell-mediated regulation of neuroinflammation involved in neurodegenerative diseases,” Journal of Neuroinflammation, 11(1), p. 201. Available at: 10.1186/s12974-014-0201-8.

Hanafy, K.A. (2013) “The role of microglia and the TLR4 pathway in neuronal apoptosis and vasospasm after subarachnoid hemorrhage,” Journal of Neuroinflammation, 10(1), p. 868. Available at: 10.1186/1742-2094-10-83.

Izzy, S. et al. (2025) “Nasal anti-CD3 monoclonal antibody ameliorates traumatic brain injury, enhances microglial phagocytosis and reduces neuroinflammation via IL-10-dependent Treg–microglia crosstalk,” Nature Neuroscience, 28(3), pp. 499–516. Available at: 10.1038/s41593-025-01877-7.

Jiang, W. et al. (2024) “CD4^+^ CD11_B_^+^ T cells infiltrate and aggravate the traumatic brain injury depending on brain-to-cervical lymph node signaling,” CNS Neuroscience & Therapeutics, 30(3), p. e14673. Available at: 10.1111/cns.14673.

Kleinschnitz, C. et al. (2010) “Early detrimental T-cell effects in experimental cerebral ischemia are neither related to adaptive immunity nor thrombus formation,” Blood, 115(18), pp. 3835–3842. Available at: 10.1182/blood-2009-10-249078.

Lauzier, D.C. et al. (2023) “Early Brain Injury After Subarachnoid Hemorrhage: Incidence and Mechanisms,” Stroke, 54(5), pp. 1426–1440. Available at: 10.1161/STROKEAHA.122.040072.

Lazarevic, I. et al. (2023) “The choroid plexus acts as an immune cell reservoir and brain entry site in experimental autoimmune encephalomyelitis,” Fluids and Barriers of the CNS, 20(1), p. 39. Available at: 10.1186/s12987-023-00441-4.

Li, Y. et al. (2025) “Single-cell sequencing reveals intracranial microvasculature-derived CXCL12 promotes CD8+ T-cell infiltration and blood–brain barrier dysfunction after subarachnoid hemorrhage in mice,” Journal of Neuroinflammation, 22(1), p. 116. Available at: 10.1186/s12974-025-03444-0.

Llovera, G. et al. (2017) “The choroid plexus is a key cerebral invasion route for T cells after stroke,” Acta Neuropathologica, 134(6), pp. 851–868. Available at: 10.1007/s00401-017-1758-y.

Macdonald, R.L. (2014) “Delayed neurological deterioration after subarachnoid haemorrhage,” Nature Reviews Neurology, 10(1), pp. 44–58. Available at: 10.1038/nrneurol.2013.246.

Macdonald, R.L. and Schweizer, T.A. (2017) “Spontaneous subarachnoid haemorrhage,” The Lancet, 389(10069), pp. 655–666. Available at: 10.1016/S0140-6736(16)30668-7.

Magatti, M. et al. (2023) “Systemic immune response in young and elderly patients after traumatic brain injury,” Immunity & Ageing, 20(1), p. 41. Available at: 10.1186/s12979-023-00369-1.

Mao, J. et al. (2024) “Microglia-derived ADAM9 promote GHRH neurons pyroptosis by Mad2L2-JNK-caspase-1 pathway in subarachnoid hemorrhage,” Journal of Neuroinflammation, 21(1), p. 302. Available at: 10.1186/s12974-024-03299-x.

Moraes, L. et al. (2020) “TH17/Treg imbalance and IL-17A increase after severe aneurysmal subarachnoid hemorrhage,” Journal of Neuroimmunology, 346, p. 577310. Available at: 10.1016/j.jneuroim.2020.577310.

Moro, F. et al. (2021) “Efficacy of acute administration of inhaled argon on traumatic brain injury in mice,” British Journal of Anaesthesia, 126(1), pp. 256–264. Available at: 10.1016/j.bja.2020.08.027.

Provencio, J.J. et al. (2011) “Depletion of Ly6G/C+ cells ameliorates delayed cerebral vasospasm in subarachnoid hemorrhage,” Journal of Neuroimmunology, 232(1–2), pp. 94–100. Available at: 10.1016/j.jneuroim.2010.10.016.

Provencio, J.J. (2013) “Inflammation in Subarachnoid Hemorrhage and Delayed Deterioration Associated with Vasospasm: A Review,” in M. Zuccarello et al. (eds.) Cerebral Vasospasm: Neurovascular Events After Subarachnoid Hemorrhage. Vienna: Springer Vienna, pp. 233–238. Available at: 10.1007/978-3-7091-1192-5_42.

Roa, J.A. et al. (2020) “Preliminary results in the analysis of the immune response after aneurysmal subarachnoid hemorrhage,” Scientific Reports, 10(1), p. 11809. Available at: 10.1038/s41598-020-68861-y.

Romoli, M. et al. (2023) “Immunological Profile of Vasospasm after Subarachnoid Hemorrhage,” International Journal of Molecular Sciences, 24(10), p. 8856. Available at: 10.3390/ijms24108856.

Sabri, M. et al. (2009) “Anterior circulation mouse model of subarachnoid hemorrhage,” Brain Research, 1295, pp. 179–185. Available at: 10.1016/j.brainres.2009.08.021.

Schallner, N. et al. (2015) “Microglia regulate blood clearance in subarachnoid hemorrhage by heme oxygenase-1,” Journal of Clinical Investigation, 125(7), pp. 2609–2625. Available at: 10.1172/JCI78443.

Schneider, U.C. et al. (2015) “Microglia inflict delayed brain injury after subarachnoid hemorrhage,” Acta Neuropathologica, 130(2), pp. 215–231. Available at: 10.1007/s00401-015-1440-1.

Sehba, F.A. et al. (2012) “The importance of early brain injury after subarachnoid hemorrhage,” Progress in Neurobiology, 97(1), pp. 14–37. Available at: 10.1016/j.pneurobio.2012.02.003.

Shi, S.X. et al. (2023) “CD4^+^ T cells aggravate hemorrhagic brain injury,” Science Advances, 9(23), p. eabq0712. Available at: 10.1126/sciadv.abq0712.

Shichita, T. et al. (2009) “Pivotal role of cerebral interleukin-17–producing γδT cells in the delayed phase of ischemic brain injury,” Nature Medicine, 15(8), pp. 946–950. Available at: 10.1038/nm.1999.

Song, M. et al. (2023) “Th17/Treg imbalance in peripheral blood from patients with intracranial aneurysm,” Journal of Neurosurgical Sciences, 67(6). Available at: 10.23736/S0390-5616.21.05567-3.

Sun, J.-Q. et al. (2024) “SIRT2 Promotes NLRP3-Mediated Microglia Pyroptosis and Neuroinflammation via FOXO3a Pathway After Subarachnoid Hemorrhage,” *Journal of Inflammation Research*, Volume 17, pp. 11679–11698. Available at: 10.2147/JIR.S487716.

Thilak, S. et al. (2024) “Diagnosis and management of subarachnoid haemorrhage,” Nature Communications, 15(1), p. 1850. Available at: 10.1038/s41467-024-46015-2.

Van Gijn, J., Kerr, R.S. and Rinkel, G.J. (2007) “Subarachnoid haemorrhage,” The Lancet, 369(9558), pp. 306–318. Available at: 10.1016/S0140-6736(07)60153-6.

Williams, G.P. et al. (2021) “CD4 T cells mediate brain inflammation and neurodegeneration in a mouse model of Parkinson’s disease,” Brain, 144(7), pp. 2047–2059. Available at: 10.1093/brain/awab103.

Xie, L. et al. (2019) “Experimental ischemic stroke induces long-term T cell activation in the brain,” Journal of Cerebral Blood Flow & Metabolism, 39(11), pp. 2268–2276. Available at: 10.1177/0271678X18792372.

Xu, H.-L. et al. (2015) “Protective role of fingolimod (FTY720) in rats subjected to subarachnoid hemorrhage,” Journal of Neuroinflammation, 12(1), p. 16. Available at: 10.1186/s12974-015-0234-7.

Yshii, L. et al. (2022) “Astrocyte-targeted gene delivery of interleukin 2 specifically increases brain-resident regulatory T cell numbers and protects against pathological neuroinflammation,” Nature Immunology, 23(6), pp. 878–891. Available at: 10.1038/s41590-022-01208-z.

Zhang, A. et al. (2023) “Clinical Potential of Immunotherapies in Subarachnoid Hemorrhage Treatment: Mechanistic Dissection of Innate and Adaptive Immune Responses,” Aging and disease, 14(5), p. 1533. Available at: 10.14336/AD.2023.0126.

